# Nutritional Protection Against Antipsychotic-Related Brain Injury: Combined Omega-3 and Vitamin E Effects on the Cerebellum

**DOI:** 10.64898/2026.01.06.697920

**Authors:** Alake Tomiwa Goodness, Olakunle James Onaolapo, Adejoke Yetunde Onaolapo

## Abstract

Haloperidol is a widely used typical antipsychotic for schizophrenia but is associated with neurotoxic effects mediated by oxidative stress, neuroinflammation, and neuronal disruption. Omega-3 fatty acids possess anti-inflammatory and neuroprotective properties, while vitamin E is a potent lipid-soluble antioxidant. Although each has shown protective effects individually, their combined efficacy against haloperidol-induced cerebellar toxicity remains poorly understood. This study investigated the neuroprotective effects of co-administered omega-3 fatty acids and vitamin E on haloperidol-induced behavioural, biochemical, and histological alterations in the rat cerebellum. Sixty male Wistar rats (80–100 g) were randomly assigned to six groups (n = 10). Control animals received a standard diet and olive oil, while treatment groups received omega-3 fatty acids (500 mg/kg of feed), vitamin E (10 IU/kg, orally), haloperidol (1 mg/kg, intraperitoneally), or their combinations for 28 days. Haloperidol-treated rats showed significant reductions in body weight, feed intake, locomotor activity, rearing, and spatial working memory, alongside increased grooming behaviour and catalepsy (p < 0.05). These behavioural deficits were markedly attenuated by omega-3 fatty acids and vitamin E, particularly when administered together. Haloperidol also increased malondialdehyde and pro-inflammatory cytokines (IL-1β, IL-6, TNF-α) while reducing total antioxidant capacity, indicating heightened oxidative stress and inflammation. Co-treatment with omega-3 fatty acids and vitamin E significantly reversed these biochemical alterations. Histological analysis revealed pronounced cerebellar neuronal degeneration in haloperidol-only rats, whereas cerebellar architecture was largely preserved in co-treated groups. Overall, combined omega-3 fatty acid and vitamin E supplementation provided significant neuroprotection against haloperidol-induced cerebellar toxicity and motor impairment, supporting their potential role as adjunct therapies in reducing antipsychotic-related neurodegeneration.

## 1. Introduction

Haloperidol is a first-generation (typical) antipsychotic widely used in the treatment of schizophrenia and other psychotic disorders. However, its clinical utility is limited by a strong association with extrapyramidal side effects and neurodegenerative-like features, particularly Parkinsonism [1–3]. By antagonising dopamine D2 receptors, haloperidol disrupts dopaminergic neurotransmission, leading to motor impairments such as tremors, rigidity, and bradykinesia—symptoms that closely resemble those observed in Parkinson’s disease [4–6]. Beyond these motor disturbances, chronic haloperidol exposure has been implicated in oxidative stress, mitochondrial dysfunction, neuroinflammation, and progressive neuronal loss, raising concerns about its potential contribution to long-term neurotoxicity [7].

In recent years, omega-3 fatty acids have attracted increasing attention as potential neuroprotective agents. Found abundantly in fish oils and certain plant sources, omega-3 fatty acids are essential polyunsaturated lipids with well-established anti-inflammatory, antioxidant, and neurorestorative properties [8–11]. Experimental evidence suggests that omega-3 fatty acids support neuronal membrane integrity, enhance synaptic plasticity, and attenuate oxidative damage—mechanisms that may counteract haloperidol-induced neurotoxic pathways [12–14]. Their ability to modulate neuroinflammatory responses and reduce reactive oxygen species further highlights their therapeutic promise.

Vitamin E, a potent lipid-soluble antioxidant, also plays a critical role in neuroprotection [15–18]. As a major chain-breaking antioxidant within cell membranes, vitamin E protects polyunsaturated fatty acids from lipid peroxidation and limits oxidative damage [17, 19]. Its neuroprotective effects have been demonstrated in several models of chemically induced neurotoxicity, where it preserves membrane structure, reduces oxidative stress, and modulates inflammatory signalling [20, 21]. When combined with omega-3 fatty acids, vitamin E may enhance membrane stability and antioxidant efficiency, suggesting a complementary protective interaction [22]. Against this background, the present study examined the neuroprotective effects of combined omega-3 fatty acids and vitamin E supplementation on haloperidol-induced alterations in rats. We assessed changes in body weight, feed intake, open-field behaviours, and memory performance, alongside biochemical markers of oxidative stress, antioxidant capacity, and pro-inflammatory cytokines, including interleukin (IL)-1β and IL-6. Histological analyses were also conducted to determine whether combined supplementation could prevent or attenuate haloperidol-induced structural damage in the cerebellum.

## 2. Materials and Methods

### 2.1 Chemicals and drugs

Haloperidol (Haldol®, Janssen Pharmaceuticals, Belgium), omega-3 fatty acid capsules containing eicosapentaenoic acid (EPA) and docosahexaenoic acid (DHA) (Pharmatec Ltd., UK), and vitamin E (α-tocopherol acetate; Nature Made®, USA), Normal saline, Assay kits for lipid peroxidation (malondialdehyde), interleukin-10, tumour necrosis factor alpha (TNF α) and total antioxidant capacity (TAC) (Biovision Inc., Milpitas, CA, USA).

### 2.2 Animals

Wistar rats weighing 80-100 g each used for this study were obtained from Empire Breeders, Osogbo, Osun State, Nigeria. Rats were housed in groups of six, in wooden cages, inside temperature-controlled quarters (22-25 degree Celsius) with 12 hours of light daily. All animals were fed commercially-available standard rodent chow (Calories: 29% protein, 13% fat, 58% carbohydrate) from weaning. Animals had access to food and water *ad-libitum*, except during the behavioural tests. All procedures were conducted in compliance with approved institutional protocols and adhered to the provisions for animal care and use outlined in the European Council Directive (EU 2010/63) on the protection of animals used for scientific purposes

### 2.3 Experimental methodology

Sixty (60) rats, each weighing 80-100g were randomly allocated into six (6) groups (n=10 per group). Group A, the control group, were fed standard rat chow and administered olive oil orally at 0.5 ml/kg body weight. Rats in group B served as omega 3 acid + vitamin E control and were given omega 3 fatty acid incorporated into rodent chow at 500 mg/kg [9] of feed and administered Vitamin E orally at 10 ml/kg [23]. Rats in group C, (Haloperidol group), were fed rodent chow and administered haloperidol intraperitoneally at 1 mg/kg body weight [24, 25] and administered olive oil orally at 0.5ml/kg body weight; Group D were given omega 3 fatty acid incorporated into rodent chow at 500 mg/kg of feed plus 1mg/kg of haloperidol and 0.5 ml of inactive oil. Group E received standard rat chow along with vitamin E administered at 10 ml/kg body weight and haloperidol at 1mg/kg. Group F received omega-3 fatty acid incorporated into rodent chow at a concentration of 500 mg/kg of feed, in addition to haloperidol (1 mg/kg) and vitamin E (10 ml/kg body weight).

Feed intake was measure daily, while body weight was measured weekly. Administration was done orally, Intraperitoneal and Dietary with the aid of a cannula for 28 days. On day 29, all animals were subjected to behavioural testing. Twenty-four hours post-testing, animals were euthanized, and blood samples were collected via cardiac puncture to assess oxidative stress and inflammatory markers, including Total Antioxidant Capacity (TAC), Malondialdehyde (MDA), Tumor Necrosis Factor-alpha (TNF-α), and interleukins (IL-6, IL-10, IL-1β). The cerebellum was sectioned, fixed, and processed for histological analysis using haematoxylin and eosin (H&E) and Cresyl Fast Violet staining.

### 2.4 Open field Novelty induced Behaviours

Open-field responses in rats assesses arousal, inhibitory, and inspective exploratory behaviours, as well as anxiety behaviours. Stereotypic behaviours, including grooming, have also been measured using this paradigm. These behaviours are typically considered fundamental and signify a rodent’s capacity for exploration. Ten minutes of behaviours in the open field, including grooming, rearing, and horizontal locomotion, were observed and recorded as previously described by [26] The open-field paradigm consisted of a square enclosure with a rigid floor, measuring 36 x 36 x 26 cm. The wood was painted white, and the floor was segmented by permanent blue markings into 16 equal squares. Horizontal locomotion (number of floor units traversed by all paws), rearing frequency (number of instances the rat stood on its hind legs, either with its forelimbs against the walls of the observation cage or freely in the air), and grooming frequency (number of body-cleaning actions involving paws, licking of the body and pubis with the mouth, and face-washing behaviours indicative of stereotypic activity) within a 10-minute period were documented as previously described [26].

### 2.5 Y maze spatial working memory

The Y-maze spatial working memory test exploits the innate tendency of rodents to explore novel environments. The apparatus consists of a wooden maze with three identical arms arranged at 120° angles to form a “Y” shape. Each arm measures approximately 15 inches in length and 3.5 inches in width, with walls about 3 inches high. Each rat was placed in one arm of the maze and allowed to explore freely, with arm entries recorded when the animal’s tail had completely entered the next arm. The sequence of arm entries was documented as previously described [27].

### 2.6 Catalepsy Bar Test

The inability to alter an enforced posture caused by muscle rigidity is commonly measured by catalepsy bar tests. Rats were carefully placed with their forelimbs on the bar and their hindlimbs on the apparatus’s floor. The bar test was adjusted to a height of 12 cm, and the time it took the rat to remove both paws from the bar was recorded using a timer. An indicator of the severity of catalepsy is the amount of time it takes the rat to straighten out this posture. A normal rat will shift its position in a matter of seconds, whereas a cataleptic rat will grip onto the bar for an extended amount of time [28, 29].

### 2.7 Estimation of MDA Content (Lipid Peroxidation)

The 2-Thiobarbituric Acid Reactive Substances (TBARS) assay quantifies lipid peroxidation by measuring MDA-TBA adduct formation at 532 nm after reacting processed plasma samples with indicator and acid reagents [27].

### 2.8 Antioxidant Status

Total antioxidant capacity was measured using the Trolox Equivalent Antioxidant Capacity Assay that is based on the ability of antioxidants within a sample to react with oxidized products [9].

### 2.9 Inflammatory Markers

Interleukin1β, IL-6, IL-10 and TNF alpha levels was assayed using enzyme-linked immunosorbent assay (ELISA) techniques with commercially available kits (Enzo Life Sciences Inc. NY, USA).

## 2.10 Tissue Histology

Rat cerebellum was processed for paraffin-embedding, cut at 5 µm and stained with haematoxylin and eosin and cresyl fast violet stains

### 2.11 Photomicrography

Histological slides of the cerebellum were examined under an Olympus binocular light microscope. Images were captured using a Canon PowerShot 2500 Digital camera.

### 2.12 Statistical Analysis

Data was analysed using Chris Rorden’s ANOVA for Windows (version 0.98). Analysis of data was by One-way analysis of variance (ANOVA) and a post-hoc test (Tukey HSD) was used. Results were expressed as mean ± S.E.M and p < 0.05 was taken as the accepted level of significant difference.

## 3. Results

### 3.1 Effect of omega 3 fatty acid and vitamin E on body weight

Figure 1 shows the effect of omega 3 fatty acid (OMEGA) and vitamin E (VIT) on relative change in body weight in haloperidol (HALO)-induced neurotoxicity in rats. There was a significant decrease in body weight with HALO and a significant increase with HALO =OMEGA, HALO+VIT, and HALO+OMEGA+VIT compared to control. Compared with HALO, body weight increased significantly with HALO+OMEGA, HALO+VIT, HALO+OMEGA+VIT. Compared with OMEGA +VIT, body weight increased with HALO+OMEGA+VIT.

**Figure 1.**
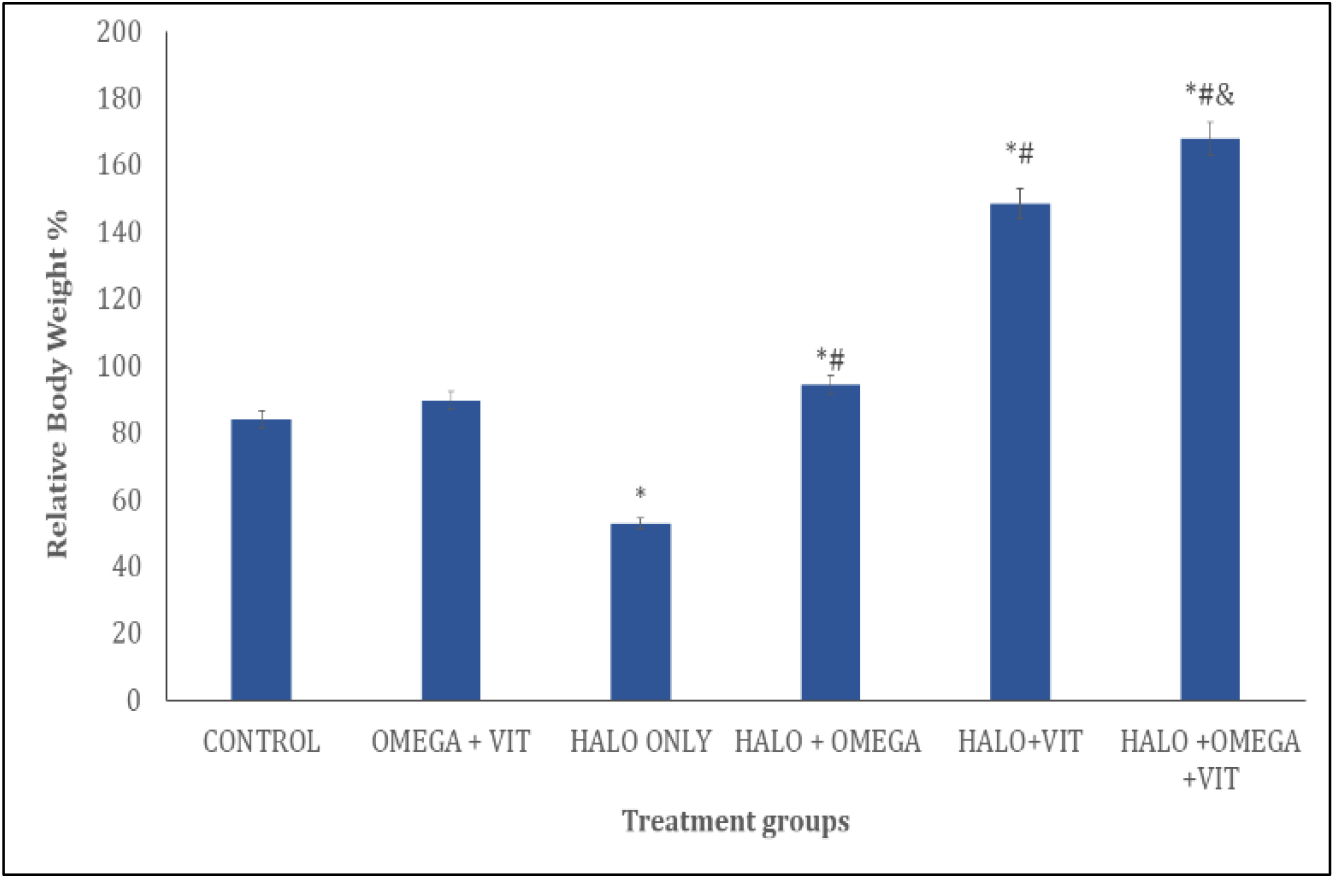
Effect of omega 3 fatty acid and vitamin E on relative change in body weight in haloperidol induced neurotoxicity in rats. Each bar represents Mean ± S.E.M, ^*^p<0.05 significant difference from control, #p<0.05 significant difference from haloperidol, ^&^p<0.005 significant difference from Omega and Vitamin E control. Numbers of rats per treatment group=10. HALO: Haloperidol, OMEGA: Omega 3 fatty acid and, VIT: Vitamin-E

### 3.2 Effect of omega 3 fatty acid and vitamin E on feed intake

Figure 2 shows the effect of omega 3 fatty acid and vitamin E on relative change in feed intake in haloperidol-induced neurotoxicity in rats. There was a significant decrease in feed intake with HALO, HALO+OMEGA and HALO+VIT and a significant increase with HALO+OMEGA+VIT compared with control. Compared with HALO, feed intake increased significantly with HALO+OMEGA, HALO+VIT, HALO+OMEGA+VIT. Compared with OMEGA +VIT, feed intake increased with HALO+OMEGA+VIT.

**Figure 2.**
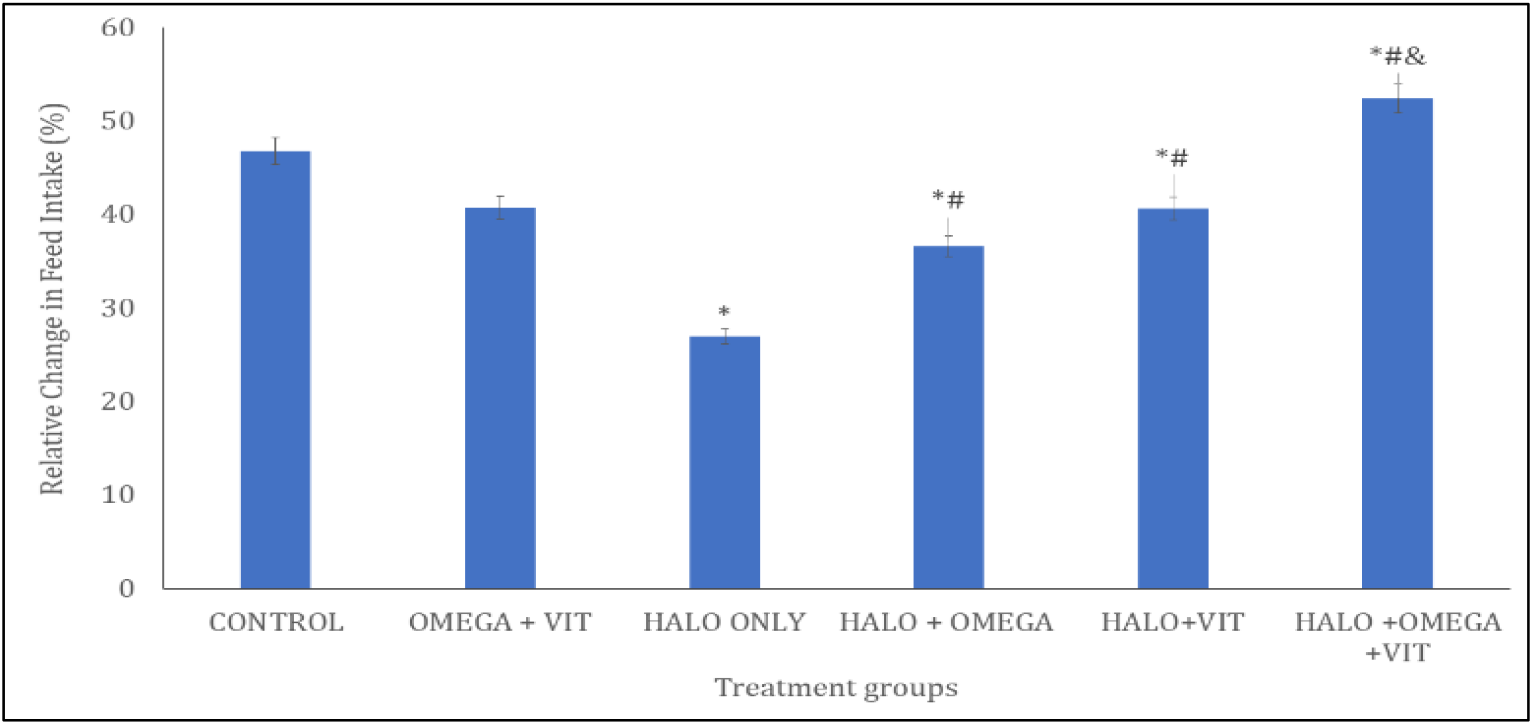
Effect of omega 3 fatty acid and vitamin E on relative change in feed intake in haloperidol induced neurotoxicity in rats. Each bar represents Mean ± S.E.M, ^*^p<0.05 significant difference from control, #p<0.05 significant difference from haloperidol, ^&^p<0.005 significant difference from Omega and Vitamin E control. Numbers of rats per treatment group=10. HALO: Haloperidol, OMEGA: Omega 3 fatty acid and, VIT: Vitamin-E

### 3.3 Effect of omega 3 fatty acid and Vitamin E on Open-field Exploratory Behaviours

Figures 3 and 4 show the effects of omega-3 fatty acid and vitamin E on horizontal locomotion and rearing activities respectively. **Figure 3** shows the effect of omega 3 fatty acid and vitamin E on horizontal locomotor activity (line crossing) in haloperidol-induced neurotoxicity in rats in the open field arena. There was a significant decrease in horizontal locomotor activity with HALO, HALO+OMEGA and, HALO+VIT compared with control. Compared with HALO, horizontal locomotion increased significantly with HALO+OMEGA, HALO+VIT, HALO+OMEGA+VIT. Compared with OMEGA +VIT, locomotor activity did not significantly differ with HALO+OMEGA+VIT.

**Figure 3.**
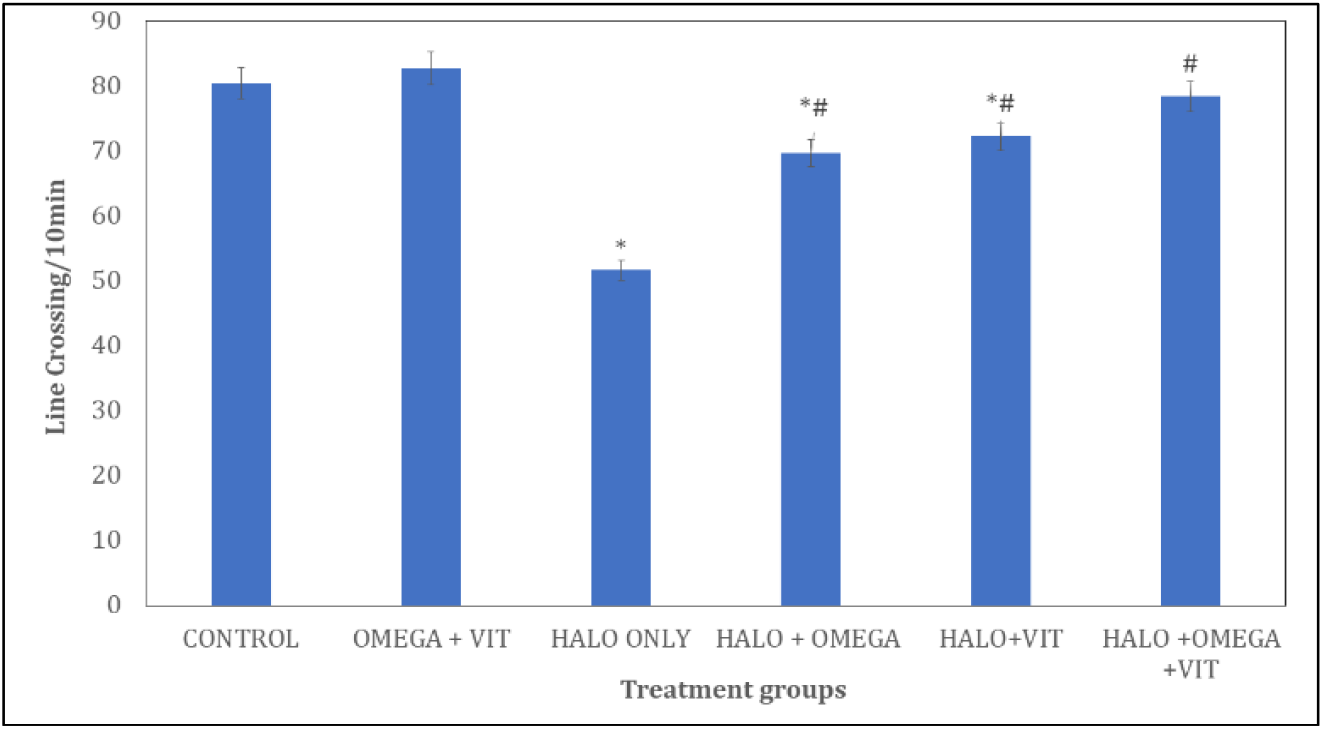
Effect of omega 3 fatty acid and vitamin E on locomotor activity in haloperidol induced neurotoxicity in rats. Each bar represents Mean ± S.E.M, ^*^p<0.05 significant difference from control, #p<0.05 significant difference from haloperidol, ^&^p<0.005 significant difference from Omega and Vitamin E control. Numbers of rats per treatment group=10. HALO: Haloperidol, OMEGA: Omega 3 fatty acid and, VIT: Vitamin-E.

**Figure 4.**
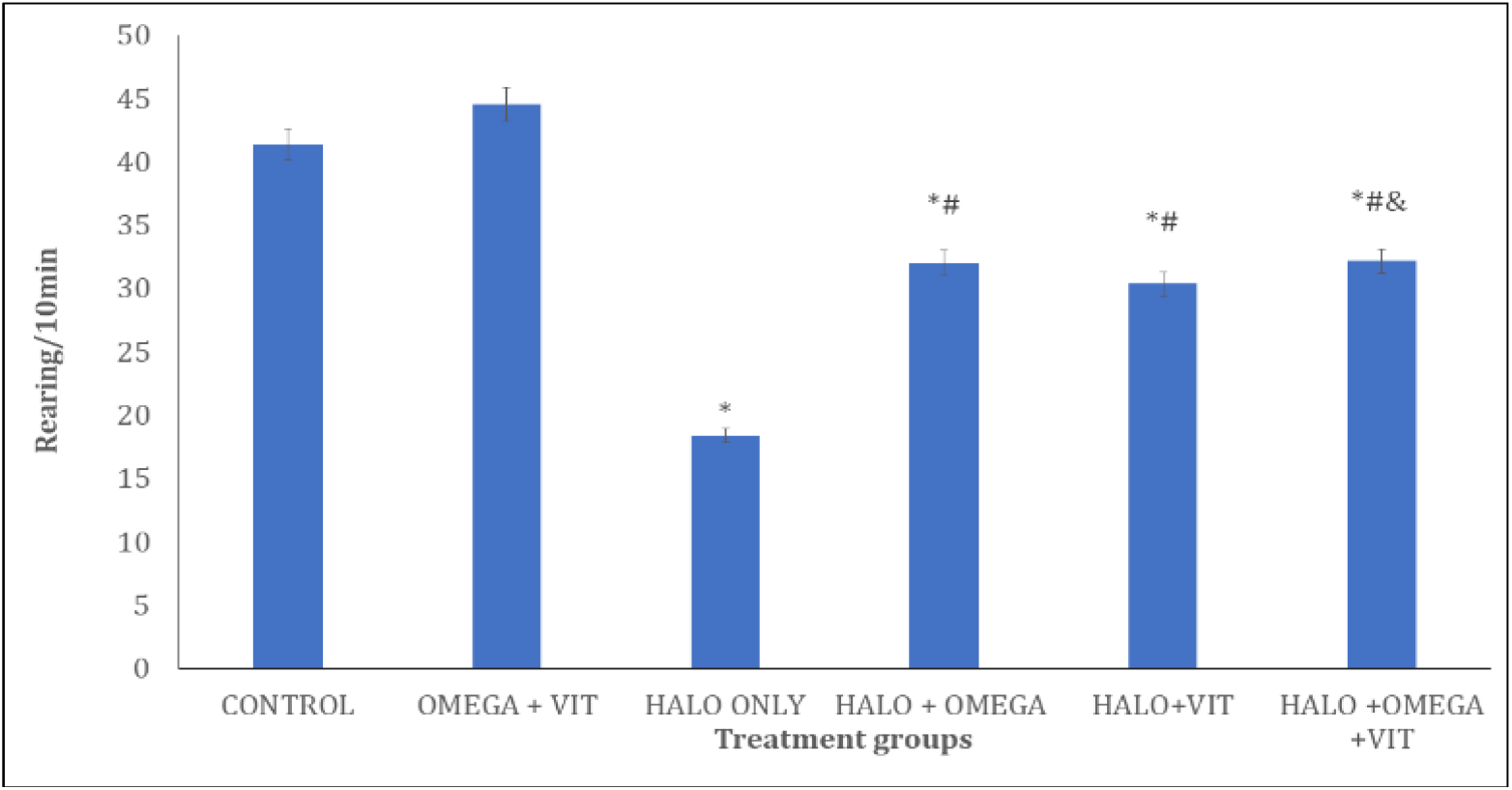
Effect of omega 3 fatty acid and vitamin E on rearing activity in haloperidol induced neurotoxicity in rats. Each bar represents Mean ± S.E.M, ^*^p<0.05 significant difference from control, #p<0.05 significant difference from haloperidol, ^&^p<0.005 significant difference from Omega and Vitamin E control. Numbers of rats per treatment group=10. HALO: Haloperidol, OMEGA: Omega 3 fatty acid and, VIT: Vitamin-E.

Figure 4 shows the effect of omega 3 fatty acid and vitamin E on vertical locomotor activity (rearing) in haloperidol-induced neurotoxicity in rats. There was a significant decrease in rearing with HALO, HALO+OMEGA, HALO+VIT and, HALO+OMEGA+VIT compared to control. Compared with HALO, rearing activity increased significantly with HALO+OMEGA, HALO+VIT, HALO+OMEGA+VIT. Compared with OMEGA+VIT, rearing decreased with HALO+OMEGA+VIT.

### 3.4 Effect of omega 3 fatty acid and Vitamin E on self-grooming behaviour

Figure 5 shows the effect of omega 3 fatty acid and vitamin E on self-grooming behaviour in haloperidol-induced neurotoxicity in rats. There was a significant increase with HALO, HALO+OMEGA and, HALO+VIT when compared with control. Compared with HALO, self-grooming behaviour decreased significantly with HALO+OMEGA, HALO+VIT, HALO+OMEGA+VIT. Compared with OMEGA +VIT, self-grooming increased with HALO+OMEGA+VIT.

**Figure 5.**
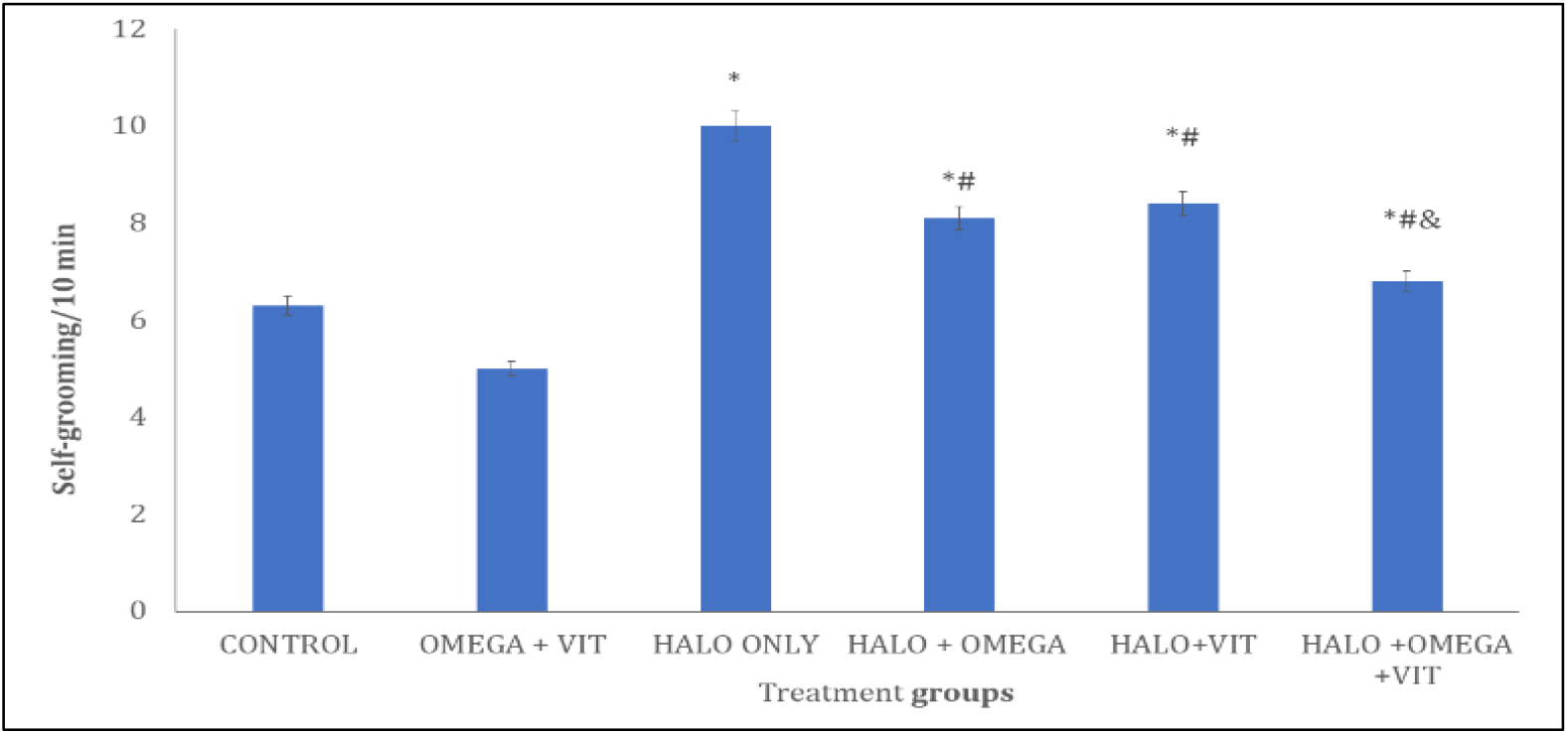
Effect of omega 3 fatty acid and vitamin E on self-grooming behaviours in haloperidol-induced neurotoxicity in rats. Each bar represents Mean ± S.E.M, ^*^p<0.05 significant difference from control, #p<0.05 significant difference from haloperidol, ^&^p<0.005 significant difference from Omega +Vitamin E control. Numbers of rats per treatment group=10. HALO: Haloperidol, OMEGA: Omega 3 fatty acid and, VIT: Vitamin-E.

### 3.5 Effect of Omega 3 fatty acid and Vitamin E on spatial working memory

Figure 6 shows the effect of omega-3 fatty acid and vit amin E on spatial working memory in the Y-maze. Spatial working memory scores measured as % alternation/5 min increased significantly (p < 0.001) in groups administered OMEGA+VIT, and decreased with HALO, HALO+OMEGA, HALO+VIT and HALO+OMEGA+VIT compared to control. Compared to HALO, memory scores increased with HALO+OMEGA, HALO+VIT, HALO+OMEGA+VIT. Compared with OMEGA +VIT, memory scores decreased with HALO+OMEGA+VIT.

**Figure 6.**
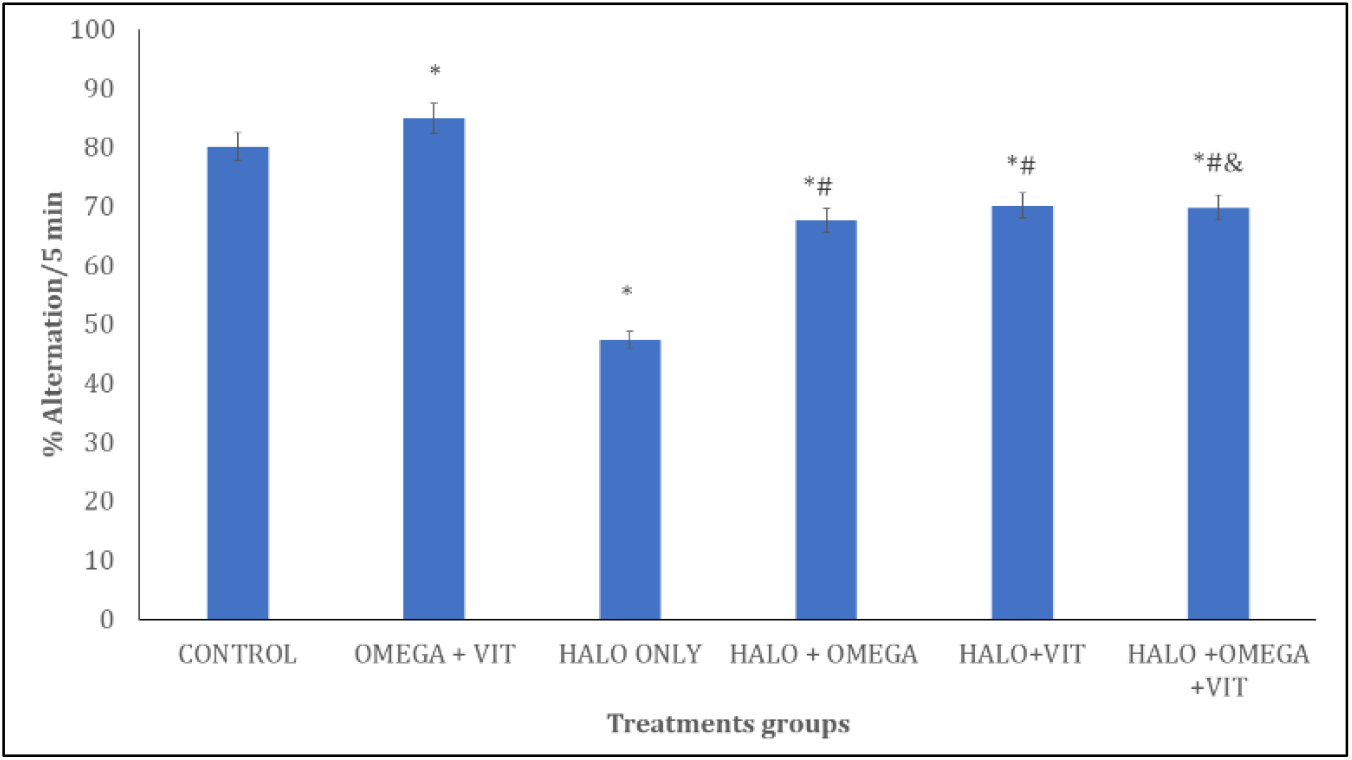
Effect of omega 3 fatty acid and vitamin E on Y-maze spatial working memory in haloperidol-induced neurotoxicity in rats. Each bar represents Mean ± S.E.M, ^*^p<0.05 significant difference from control, #p<0.05 significant difference from haloperidol, ^&^p<0.005 significant difference from Omega +Vitamin E control. Numbers of rats per treatment group=10. HALO: Haloperidol, OMEGA: Omega 3 fatty acid and, VIT: Vitamin-E.

### 3.6 Effect of Omega 3 fatty acid and Vitamin E on Catalepsy

Figure 7 shows the effect of omega 3 fatty acid and vitamin E on Immobility time in the catalepsy test. There was a significant increase in mobility time with HALO, HALO+OMEGA and, HALO+VIT and a decrease with HALO+OMEGA+VIT compared with control. Compared with HALO, immobility time decreased with HALO+OMEGA, HALO+VIT and HALO+OMEGA+VIT. Compared with OMEGA+VIT, immobility time decreased with HALO+OMEGA+VIT.

**Figure 7.**
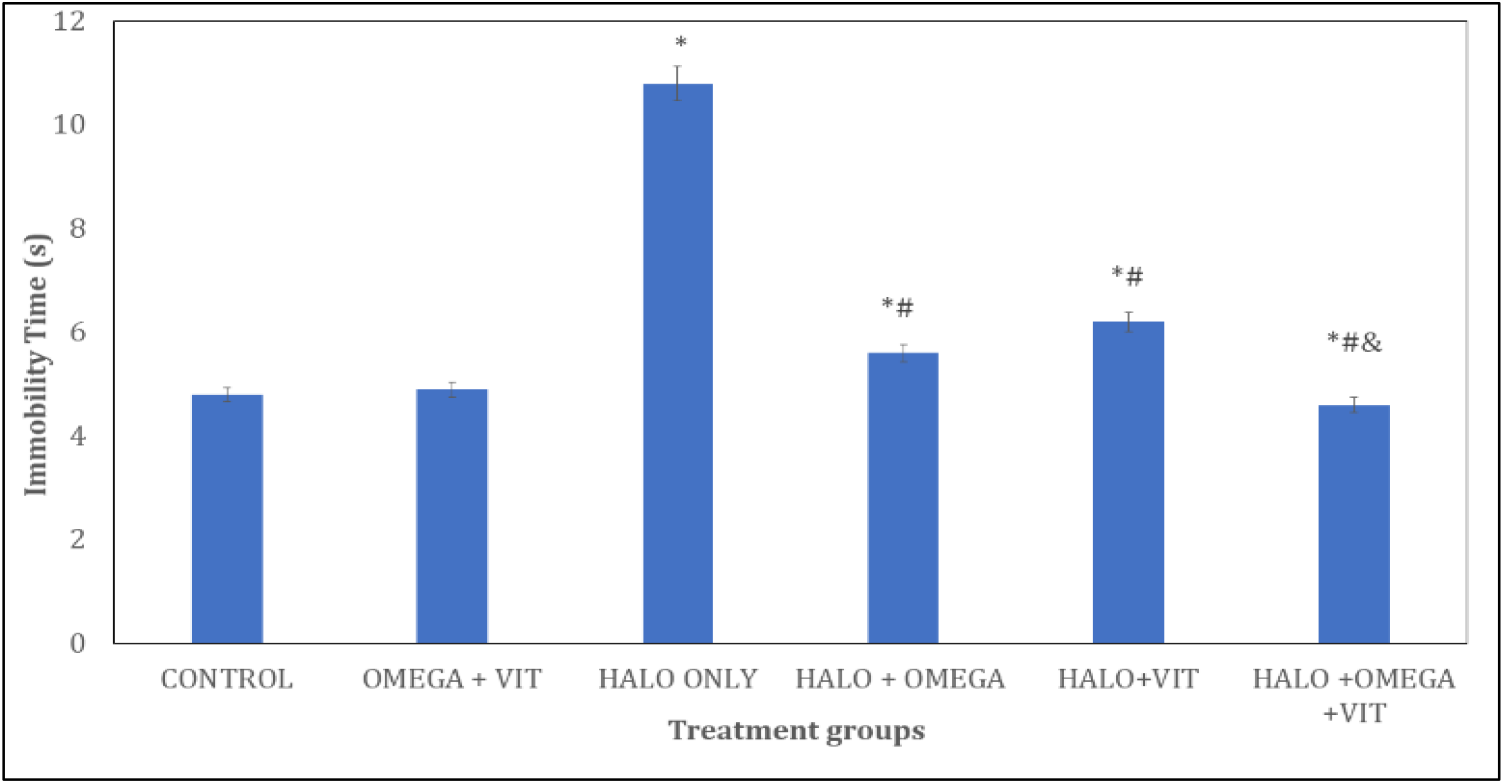
Effect of omega 3 fatty acid and vitamin E on immobility time in the catalepsy bar test. Each bar represents Mean ± S.E.M, ^*^p<0.05 significant difference from control, #p<0.05 significant difference from haloperidol, ^&^p<0.005 significant difference from Omega +Vitamin E control. Numbers of rats per treatment group=10. HALO: Haloperidol, OMEGA: Omega 3 fatty acid and, VIT: Vitamin-E.

### 3.7 Effect of omega 3 fatty acid and Vitamin E on oxidative stress parameters and inflammatory cytokines

Table 1 shows the effect of omega 3 fatty acid and vitamin E on Total antioxidant capacity (TAC) and lipid peroxidation, measured as malondialdehyde (MDA). Total antioxidant capacity (TAC) significantly decreased with HALO and increased with OMEGA+VIT, HALO+OMEGA, HALO+VIT and, HALO+OMEGA+VIT when compared to control. When compared with HALO, there was significant increase with HALO+OMEGA, HALO+VIT and HALO+OMEGA+VIT. Lipid peroxidation measured as malondialdehyde (MDA) significantly increased with HALO when compared to control. When compared with HALO, there was a significant decrease with HALO+OMEGA, HALO+VIT and HALO+OMEGA+VIT. Compared with OMEGA+VIT, TAC levels decreased with HALO+OMEGA+VIT.

**Table 1.** Effect of Omega3 fatty acid and Vitamin E on oxidative stress markers.

| Groups | TAC (mM TE) | MDA (μm) |
| --- | --- | --- |
| CONTROL | 6.48 ± 0.19 | 0.73 ± 0.02 |
| OMEGA+VIT | 20.39 ± 0.9<sup>*</sup> | 0.83 ± 0.01 |
| C (HALO) | 2.94 ± 0.48<sup>*</sup> | 12.23 ± 0.26<sup>*</sup> |
| HALO+OMEGA | 13.92 ± 0.12<sup>*#</sup> | 0.68 ± 0.02<sup>#</sup> |
| HALO+VIT | 12.52 ± 0.07<sup>*#</sup> | 0.77 ± 0.01<sup>#</sup> |
| HALO+OMEGA+VIT | 12.37 ± 0.1<sup>*#&</sup> | 0.74 ± 0.01<sup>#</sup> |

Data presented as Mean ± S.E.M, \*p < 0.05 significant difference from A, #p < 0.05 significant difference from D, number of rats per treatment group =10. MDA: Malondialdehyde, TAC: Total antioxidant capacity, VIT: Vitamin E, OMEGA: Omega-3 fatty acid

Table 2 shows the effect of omega 3 fatty acid and vitamin E on Interleukin (IL) -1β, 10, 6 and Tumour necrosis factor-alpha (TNF-α). Interleukin-1β levels increased significantly (p < 0.05) with HALO and decreased with HALO+OMEGA, HALO+VIT and HALO+OMEGA+VIT compared with control. Compared with HALO, IL-Iβ decreased with HALO+OMEGA, HALO+VIT and HALO+OMEGA+VIT. Compared with OMEGA+VIT, IL-Iβ decreased with HALO+OMEGA+VIT. Interleukin-10 levels increased with OMEGA+VIT, HALO +OMEGA+VIT and decreased with HALO compared with control. Compared with HALO, IL-10 increased with HALO+OMEGA, HALO+VIT and HALO+OMEGA+VIT. Compared with OMEGA+VIT, IL-10 levels increased with HALO+OMEGA+VIT.

Tumour necrosis factor-alpha levels increased significantly with OMEGA+VIT, HALO and HALO+OMEGA+VIT compared with control. Compared with HALO, TNF-α decreased with HALO+OMEGA and HALO+VIT, Compared with OMEGA+VIT, TNF-α increased with HALO+OMEGA+VIT. Interleukin-6 levels increased significantly with HALO and decreased with OMEGA+VIT and HALO+OMEGA+VIT compared with control. Compared with HALO, IL-6 decreased with HALO+OMEGA, HALO+VIT and HALO+OMEGA+VIT.

**Table 2.** Effect of Omega-3 fatty acids and Vitamin E on inflammatory cytokines.

| Groups | IL-1β (pg/ml) | IL-10 (pg/ml) | TNF (pg/ml) | IL-6 (pg/ml) |
| --- | --- | --- | --- | --- |
| A | 4.57 ± 0.64 | 6.28 ± 0.05 | 11.73 ± 0.3 | 8.33 ± 0.29 |
| B | 5.04 ± 0.02 | 12.39 ± 0.05<sup>*</sup> | 16.99 ± 1.74<sup>*</sup> | 4.68 ± 1.41<sup>*</sup> |
| C | 12.58 ± 0.04<sup>*</sup> | 2.86 ± 0.04<sup>*</sup> | 26.11 ± 1.87<sup>*</sup> | 13.89 ± 0.34<sup>*</sup> |
| D | 2.9 ± 0.32<sup>*#</sup> | 6.4 ± 0.5<sup>#</sup> | 12.29 ± 0.72<sup>#</sup> | 6.57 ± 0.66<sup>#</sup> |
| E | 2.36 ± 0.11<sup>*#</sup> | 7.95 ± 0.4<sup>#</sup> | 13.6 ± 0.75<sup>#</sup> | 7.03 ± 1.04<sup>#</sup> |
| F | 1.37 ± 0.13<sup>*#</sup> | 17.65 ± 1.19<sup>*#&</sup> | 26.55 ± 1.21<sup>#&</sup> | 2.03 ± 0.15<sup>*#</sup> |

Data presented as Mean ± S.E.M, \*p < 0.05 significant difference from A, #p < 0.05 significant difference from D, number of rats per treatment group =10. A: Control B: OMEGA+VIT, C: HALO, D: HALO+OMEGA, E: HALO+VIT F: HALO+OMEGA+VIT. TNF: Tumour necrosis factor -α, IL: Interleukin. VIT: Vitamin E, OMEGA: Omega-3 fatty acid, HALO: Haloperidol,

### 3.8 Effect of omega 3 fatty acid and Vitamin E on the Cerebellar cortex histomorphology

Figure 8 (A-F) and 9 (A-F) show representative photomicrographs of haematoxylin and eosin (H&E) and cresyl fast violet (CFV) stained sections of the rat cerebellar cortex respectively. Examination of the H and E-stained slides of the rats in the control (8A) and OMEGA+VIT (8B) groups revealed characteristic, well-delineated layers of the cerebellum with molecular layer, Purkinje layer and granular cell layer. Scattered within the neuropil are multipolar shaped Purkinje cells with large vesicular nucleus, granule neurons with large open-faced nuclei and scanty cytoplasm and small sized neuroglial cells in rats administered vehicle. The neuropil, which is pink staining in the H&E slides is well preserved in the CFV stained slides (9A,9B). The staining revealed the characteristic layered arrangement of the cerebellum with well-delineated multipolar Purkinje cells (PC), deeply stained granule cells (GC), neuroglia (NG), and prominent Nissl bodies in (9A, 9B). However, in the HALO group (8C, 9C), Purkinje neurons showed clear degenerative changes, including shrunken cell bodies, pale or fragmented nuclei, and a marked reduction in Nissl substance. The granular layer also appeared less densely stained, with scattered pyknotic granule cells. In the HALO+ OMEGA (8D,9D) and HALO+VIT (8E, 9E), the cerebellar cortex showed better preservation of Purkinje cells and Nissl bodies, indicating protection against neuronal injury, and this improvement was more pronounced in the HALO+OMEGA+ VIT group (8F, 9F).

**Figure 8.**
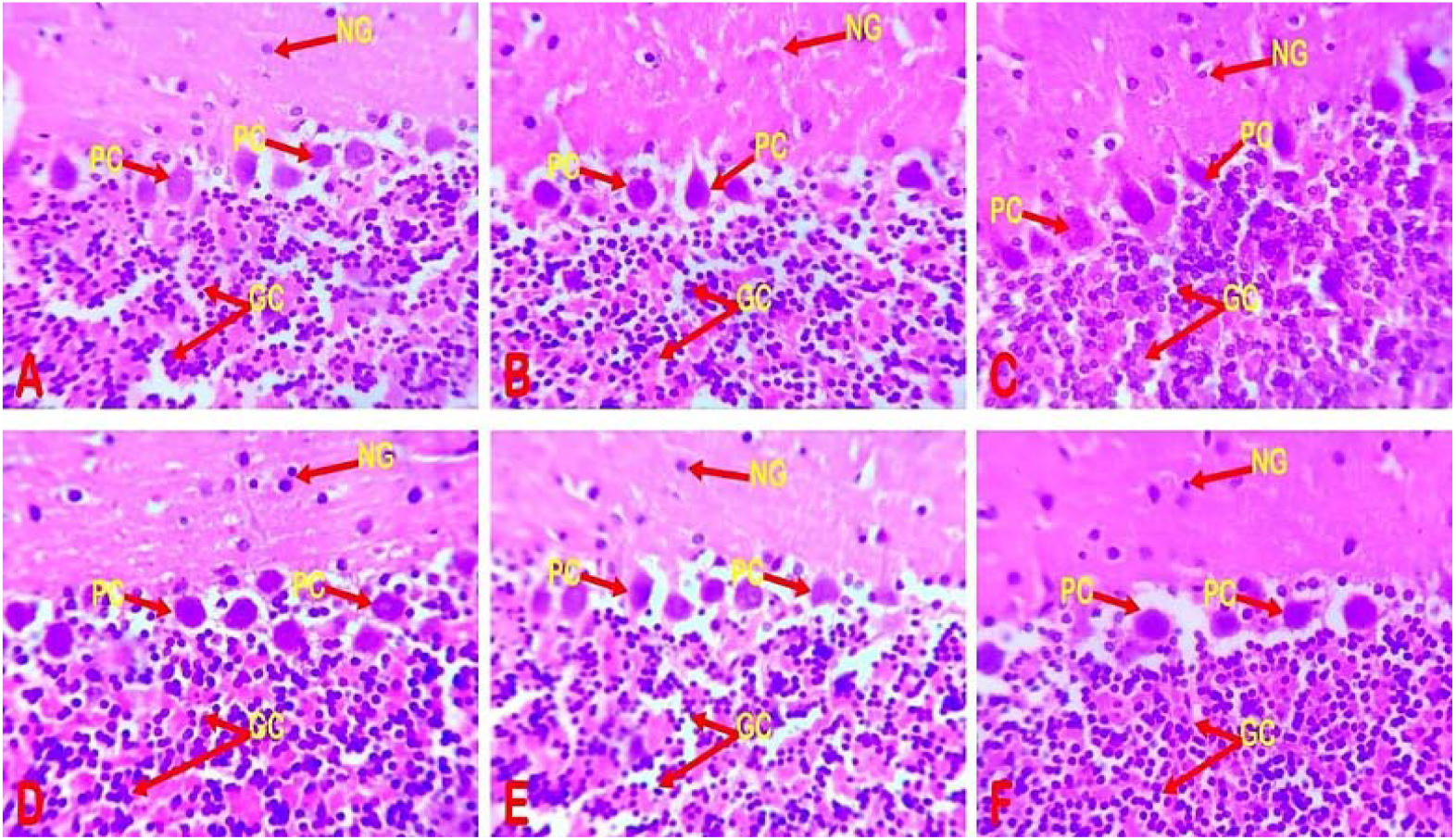
Photomicrograph of haematoxylin and eosin-stained cerebellar cortex in A: Control B: OMEGA+VIT, C: HALO, D: HALO+OMEGA, E: HALO+VIT F: HALO+OMEGA+VIT. Showing granule cells (GC) in the granule cell layers (GL), Purkinje cells (PC) in the Purkinje layers (PL) and molecular layers (ML) (Mag. X400).

**Figure 9.**
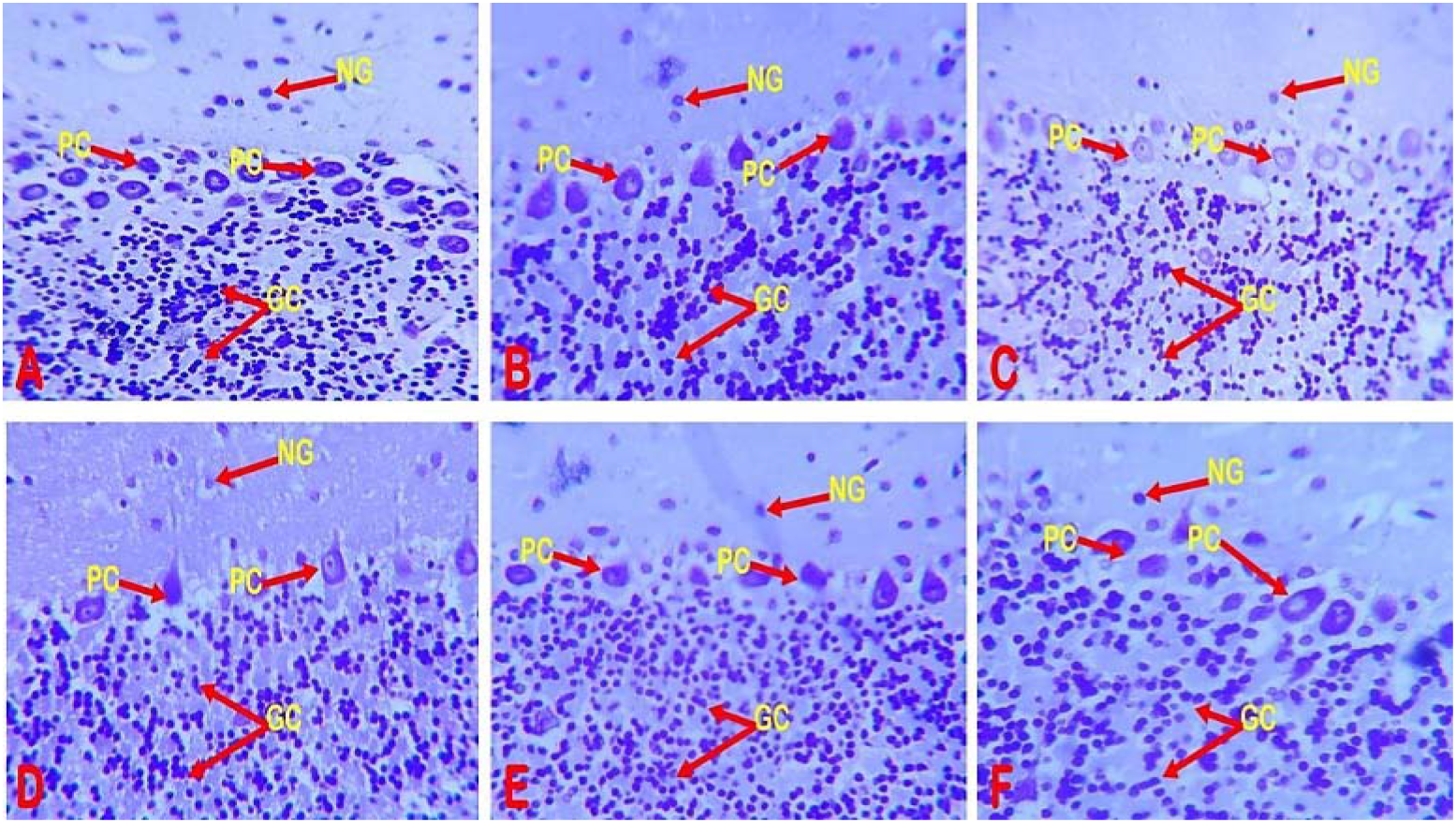
Photomicrograph of Cresyl fast violet-stained cerebellar cortex in A: Control B: OMEGA+VIT, C: HALO, D: HALO+OMEGA, E: HALO+VIT F: HALO+OMEGA+VIT. Showing granule cells (GC) in the granule cell layers (GL), Purkinje cells (PC) in the Purkinje layers (PL) and molecular layers (ML) (Mag. X400).

## Discussion

This study evaluated the neuroprotective effects of omega-3 fatty acids and vitamin E, administered individually and in combination, against haloperidol-induced neurotoxicity in rats. Across behavioural, biochemical, and histomorphological endpoints, chronic haloperidol exposure produced marked oxidative stress, neuroinflammation, motor dysfunction, cognitive impairment, and cerebellar degeneration. Supplementation with omega-3 fatty acids and vitamin E significantly mitigated these effects, with combined treatment consistently providing the most robust protection, suggesting a synergistic interaction between the two nutrients. Haloperidol significantly reduced body weight, an effect consistent with previous reports linking typical antipsychotics to metabolic suppression, anorexia, and impaired dopaminergic signalling [30,31]. Both omega-3 fatty acids and vitamin E independently improved weight outcomes, in line with studies demonstrating their roles in regulating feed intake, metabolic balance, and systemic inflammatory tone [9,15,32,33]. The greater improvement observed with combined supplementation suggests reversal of haloperidol-induced metabolic dysregulation, likely through enhanced antioxidant and anti-inflammatory capacity.

Marked reductions in locomotion, rearing, and exploratory behaviour following haloperidol treatment reflected dopamine D2 receptor blockade and extrapyramidal dysfunction [34,35]. These findings align with earlier studies reporting reduced open-field activity following dopamine antagonism [9,34]. Supplementation significantly attenuated these deficits. Omega-3 fatty acids support synaptic plasticity and dopaminergic modulation, while vitamin E preserves neuronal membrane integrity by limiting oxidative injury [38–41]. Although combined treatment restored motor function more effectively than either supplement alone, activity levels did not fully reach baseline, indicating partial but meaningful recovery. Self-grooming behaviour was elevated in haloperidol-treated rats, suggesting increased stress, anxiety, or stereotypy. While some studies report reduced grooming with haloperidol [27], increased grooming has been observed following intrastriatal haloperidol administration and under conditions of environmental novelty [42,43]. Both omega-3 fatty acids and vitamin E reduced grooming behaviour, supporting their anxiolytic and neuromodulatory effects [44–46], although grooming was not completely normalised in the combined treatment group.

Haloperidol-induced working memory impairment in the Y-maze was consistent with disrupted prefrontal dopaminergic tone and oxidative damage [9,47,48]. Omega-3 fatty acids and vitamin E improved alternation performance, with the combined group showing the greatest enhancement, consistent with previous reports [46,49,50]. These effects likely reflect improved hippocampal and cortical neuroplasticity, membrane fluidity, and synaptic preservation [51,52]. Catalepsy, a hallmark of extrapyramidal dysfunction [53,54], was significantly reduced by both supplements, with the greatest reduction observed following combined treatment. This finding reflects improved dopaminergic signalling and reduced oxidative injury within the basal ganglia, consistent with earlier studies on omega-3 fatty acids and vitamin E modulation of nigrostriatal pathways.

Biochemically, haloperidol reduced total antioxidant capacity while increasing malondialdehyde levels, confirming heightened oxidative stress and lipid peroxidation [9,53,54]. Supplementation restored antioxidant balance and reduced lipid peroxidation, with combined treatment producing the strongest effects. Omega-3 fatty acids enhance endogenous antioxidant enzyme activity, while vitamin E directly scavenges lipid radicals, explaining their complementary protective actions [38–41].

Similarly, haloperidol elevated pro-inflammatory cytokines (IL-1β, IL-6, TNF-α) and reduced IL-10, reflecting neuroinflammatory activation [55,56]. These changes were most effectively reversed by combined supplementation, consistent with the anti-inflammatory actions of omega-3-derived mediators and vitamin E inhibition of NF-κB signalling [57,58].

Histological analysis revealed pronounced cerebellar neurodegeneration following haloperidol treatment, including Purkinje cell loss, Nissl substance depletion, and cortical disorganisation. Given the cerebellum’s vulnerability to oxidative injury and its role in motor coordination, these changes provide a structural basis for the observed behavioural deficits. While individual supplementation reduced neuronal damage, combined omega-3 fatty acids and vitamin E preserved cerebellar architecture to near-control levels, highlighting the anatomical correlate of their neuroprotective effects.

## Conclusion

Chronic haloperidol administration induced significant behavioural impairments, oxidative stress, neuroinflammation, and cerebellar neurodegeneration. Supplementation with omega-3 fatty acids and vitamin E, particularly in combination, effectively mitigated these adverse effects. The superior efficacy of combined treatment reflects synergistic antioxidant, anti-inflammatory, membrane-stabilising, and neurostructural protective mechanisms. These findings support the potential use of omega-3 fatty acids and vitamin E as adjunct therapies to reduce antipsychotic-induced neurotoxicity and preserve neural integrity during long-term antipsychotic treatment.

## Funding

None

## Ethical Approval

Ethical approval for this study was obtained from faculty of basic medical sciences Ladoke Akintola university of technology, Ogbomoso (ERC Approval Number: ERCFBMSLAUTECH:112/06/2025)

## Availability of data and materials

Data generated during and analysed during the course of this study are available from the corresponding author on request.

## Notes

### Competing Interest Statement

The authors have declared no competing interest.

